# Assembling a fully-dated complete tree of life

**DOI:** 10.64898/2026.03.05.709771

**Authors:** Jonathan D. Duke, Jialiang Guo, Félix Forest, Rikki Gumbs, Emily Jane McTavish, James Rosindell

## Abstract

Time-scaled phylogenetic trees summarising evolutionary relationships are fundamental to many analyses in biology, from diversification rate estimation to conservation prioritisation. The most comprehensive available summary of these relationships, the Open Tree of Life, synthesises information from over two thousand studies into a supertree covering the full range of global biodiversity, but its use in downstream analyses is limited by the lack of divergence times. Previous work has mapped dates from Open Tree’s database of trees to certain nodes in the supertree, but for the majority of nodes no date is available. While algorithms exist to interpolate missing dates in a tree, we found that their time and memory requirements scaled quadratically with the number of nodes, which made it computationally infeasible to run them on the entire tree. In this work, we describe novel date interpolation algorithms that scale linearly with the number of nodes. These enabled us to produce a distribution of fully-dated trees containing 2.3 million extant described species, greatly expanding the scope of feasible phylogenetic analyses. We illustrate the utility of these trees by computing the most robust estimate yet of the phylogenetic diversity of the complete tree of life, incorporating both topological and temporal uncertainty.

## Introduction

The tree of life has been an object of study for centuries [1, 2], telling the story of how the diverse array of species on Earth came into being. The advent of genomic sequencing has provided vast datasets [3, 4], which have been analyzed to map out ever-larger clades of the tree [5–7]. Many fundamental questions in biology require knowledge of the evolutionary history presented in the tree. For example, speciation, extinction, and the evolution of traits are all studied using a dated tree [8–10]. Ecological analyses must account for the shared ancestry of the species they examine [11–15]. Missing data in trait databases can be imputed using phylogenetic correlations [16–18]. And in conservation, the tree-based phylogenetic diversity (PD) measure has found widespread use [19, 20].

A complete understanding of evolution requires a complete tree. Direct estimation of the entire tree from genomic data has not yet been achieved, owing both to insufficient data and to the computational infeasibility of fitting phylogenetic statistical models on such a scale. Rather than infer the entire tree using a single model, the Open Tree of Life project instead synthesises a tree by collating and summarising individual studies [21, 22], producing the most complete tree of life available today. The Open Tree synthesis algorithm stitches together over 2,000 trees from a curated database to construct a backbone tree of 150,000 species [23, 24]. This phylogenetic synthesis is combined with their taxonomic database [25, 26], producing a “supertree” that covers 2.3 million described species.

The Open Tree team make regular updates to the tree incorporating new information, improving phylogenetic coverage over time. At present, however, substantial parts of the tree come primarily from taxonomic data, resulting in extensive polytomies [27] that can produce biases in downstream analyses [28, 29]. While the true fully-resolved topology is unknown and the Open Tree synthesis algorithm explicitly does not attempt to estimate it [24], for many analyses the varying degrees of topological uncertainty across the tree could be more transparently reflected using trees sampled from a set of plausible topologies to produce distributions of possible solutions [28, 30, 31].

Crucially, the Open Tree also lacks divergence times, i.e. the estimated historical dates of the tree’s nodes, which represent common ancestors or speciation events. The lengths of the branches between nodes in units of time are used to compute phylogenetic correlations, phylogenetic diversity, and to associate evolutionary events with geological or climatic changes over the history of Earth. Previous work has mapped dates from Open Tree’s database of underlying phylogenetic trees to nodes in the supertree [6, 32, 33], but the database includes dates for only 77,000 internal nodes. Work is ongoing to improve coverage by incorporating more published dated trees - where they are available electronically - into the Open Tree database, but for the majority of nodes in the complete tree, no dates have yet been estimated.

Dating the tree therefore requires interpolation between dated nodes on a sparsely-dated tree with no branch lengths. Such interpolation methods exist [34, 35], but in their present implementations, the computational time and memory required to run them scale with the square of the number of nodes in the tree [34]. This makes interpolation across the millions of nodes in the entire tree of life infeasible. Methods that scale linearly with the number of nodes have been developed for applications such as smoothing molecular substitution rates across a tree with branch lengths in units of molecular substitutions [36, 37], but to our knowledge the dynamic programming techniques used by these algorithms have not been applied to interpolating dates across topology-only trees.

In this work, we describe stochastic algorithms that resolve the polytomies in the Open Tree by placing taxa with only taxonomic information into plausible locations in the phylogenetic backbone, automating the methods of commonly-used tree-completion tools [e.g., 38, 39, 40]. We use our algorithms to produce a distribution of bifurcating trees covering all extant described species. We then compare five linear-time methods for interpolating dates across any tree, and apply them to our distribution of bifurcating trees, resulting in a set of fully-dated complete trees of life.

## Results

### Fully-dated complete trees

Our date interpolation algorithms ran in times that scaled linearly with the number of tree nodes (**Fig 1**), which made it computationally feasible for us to produce bifurcating fully-dated trees at the scale of the entire tree of life. For example, interpolating dates across a tree of 2.3 million species using an “equal splits” interpolation method took 26 seconds (**Fig 1**). In comparison, the same method as implemented in the BLADJ algorithm in the software Phylocom scaled in both run time and memory requirement with the square of the number of nodes (**Fig 1**). We were unable to run it successfully on a tree of 2.3 million species owing to a memory requirement of 50 terabytes.

**Fig 1.**
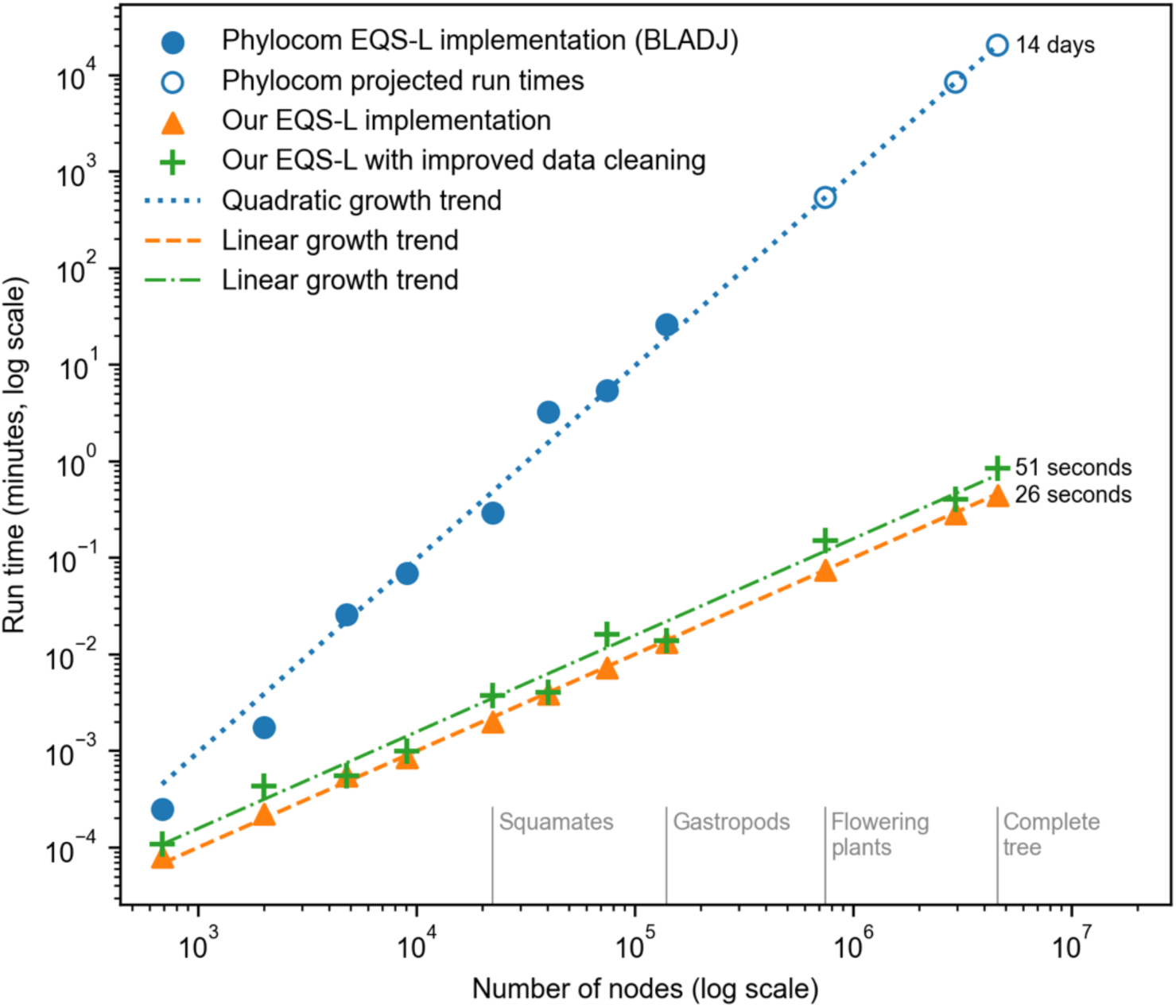
Run times for interpolation algorithms that space dates evenly along undated paths between input dates. The time taken in minutes to interpolate dates across a tree is plotted against the number of nodes in that tree, for (blue circles) the BLADJ implementation in Phylocom; (orange triangles) our implementation of the same algorithm, which we call EQS-L; (green crosses) our implementation including our improved data cleaning algorithm to make the input dates time-consistent. Each triplet of points at a given number of nodes represents the same subtree extracted from a partially-dated complete tree. The blue dotted line illustrates a growth trend in run times increasing with the square of the number of nodes, while the orange dashed line and green dash-dot line increase linearly with the number of nodes. The three blue open circles show hypothetical run times imputed from the trend, because Phylocom’s BLADJ implementation did not run successfully on trees of such size - on the complete tree, our projected run time is 14 days, but in practice the run failed owing to a memory requirement of 50 terabytes.

### Achieving temporal consistency of input dates

Our interpolation used calibration dates summarised from 334 source trees. The source trees differed from each other and from the supertree in their calibration dates, dating methodologies and topologies, so when these dates were mapped to the supertree they were not time-consistent. That is, some nodes were assigned dates younger than the date assigned to a descendant node or older than an ancestor. The BLADJ implementation includes a data cleaning step that ensures time consistency by traversing the tree from root to leaves and removing the date from any node that has a younger ancestor. In our tree, this step removed 9,121 of the 76,953 dates. Our alternative algorithm (see Materials and methods) removes only the minimum number of dates necessary to achieve time consistency across the tree. It removed only 5,000 dates, retaining more of the original information while still operating with linear time complexity (**Fig 1**).

### Comparison of interpolation algorithms

We compared five approaches for interpolating missing dates on sparsely-dated trees with no branch lengths. We tested the *equal splits* algorithm, which spaces nodes evenly along undated paths between input dates; we gave this the abbreviation EQS, after Kuhn et al. [28]. Sometimes, the algorithm must interpolate along several paths from the same start node, all of which end in a leaf node. In this case, the choice of which path is interpolated first will affect the result (see Materials and methods for further explanation and details of all approaches). We therefore tested two variants of the EQS approach: one which interpolates first along the longest available path (we call this EQS-L; this is the approach used by the BLADJ algorithm in the Phylocom software) and one which interpolates first along the shortest path (EQS-S). Our third approach used a linear combination of the dates obtained from these methods, with the parameter choice of 0.25*EQS-L + 0.75*EQS-S determined by testing the algorithms on small fully-dated trees. We called this linear combination EQS-LS. We also considered two approaches that assign dates using the expected branch lengths from a birth model with exponentially-distributed waiting times: the *log-N* (LnN) approach, which works down each lineage in a clade independently and is deterministic, and an approach we called *birth model* (BM), which randomly chooses the order of speciation events across an undated clade.

To compare the algorithms, we first used four fully-dated trees that had been produced using molecular data for every included species (see Materials and methods for details). In each tree, we removed some of the input dates at random and applied the interpolation algorithms in turn to estimate dates for the undated nodes. We then computed the date reproduction error. We repeated this analysis for seven clades of our fully-resolved complete tree, each of which had a crown age in our input data and between 19% and 73% of internal nodes dated. For all trees, we also computed mean and median pendant edge lengths, to examine how closely the algorithms maintained the distribution of branch lengths in the tree.

We found that, in trees where fewer than 70% of nodes needed to be interpolated, the five interpolation methods all performed similarly, with an average absolute reproduction error of 1.5-3% of the crown age (**Fig 2** row 1). For trees where more than 70% of nodes required an interpolated date, the EQS-LS approach produced the lowest reproduction error of 3-4%, while the error for the other approaches increased more rapidly with the degree of interpolation, especially when interpolating more than 95% of dates.

**Fig 2:**
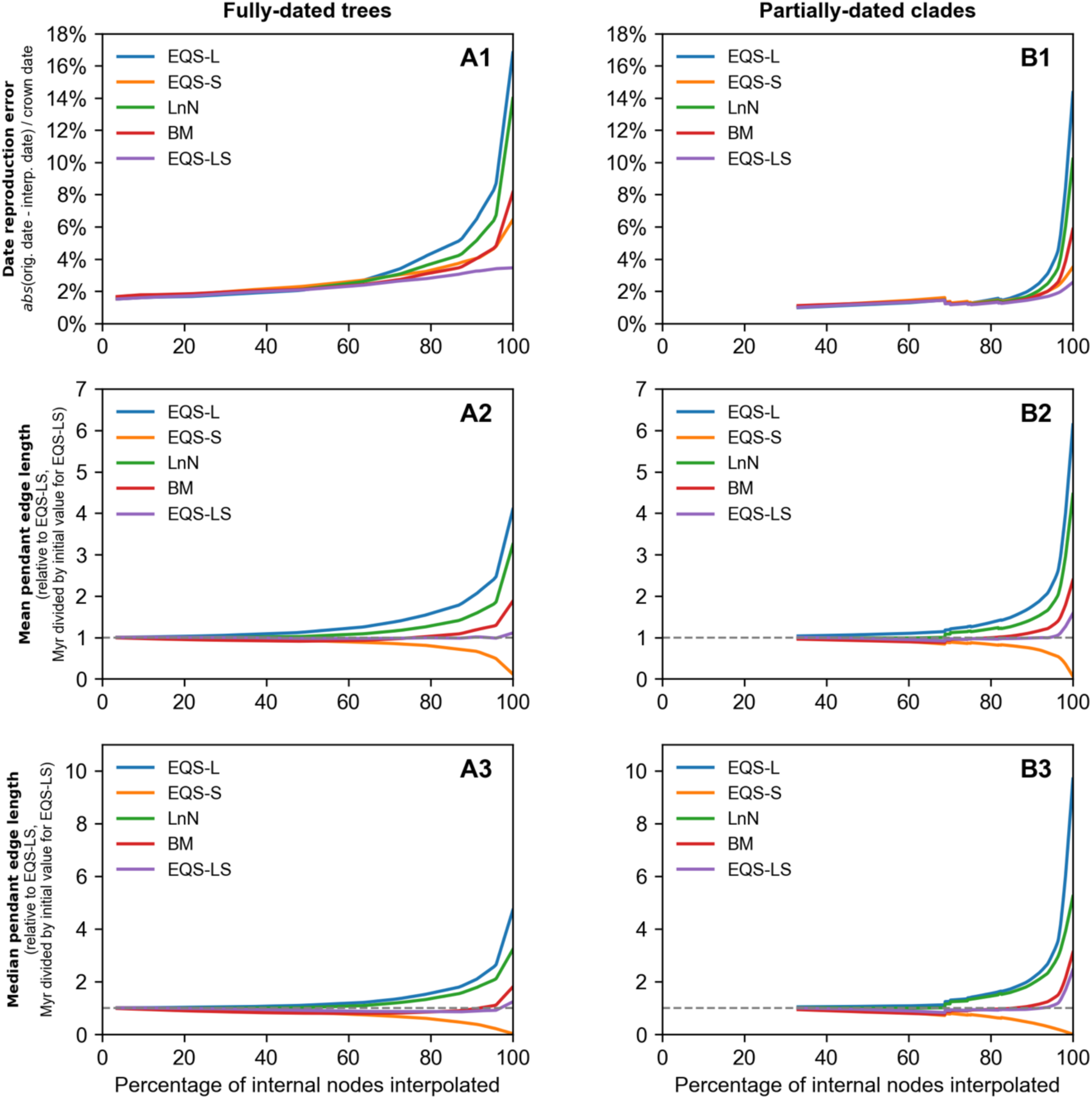
Performance of date interpolation algorithms. Row 1: the average error in date reproduction tests across (column A) four fully-dated phylogenetic trees and (column B) seven subtrees of one sample tree in our distribution of fully-resolved complete trees of life. We removed a proportion of calibration dates at random from the tree, then interpolated dates across all undated internal nodes, and compared the interpolated dates with the original dates we had removed. The full datasets are supplied in the supplementary materials. Row 2: mean pendant edge length in the tree, relative to (for column A) the mean pendant edge length in the fully-dated tree, or (for column B) the mean pendant edge length in the subtree when using all available calibration dates and missing dates interpolated using the EQS-LS algorithm. Row 3: as row 2, but for median pendant edge lengths rather than mean.

The EQS-LS algorithm also kept the distribution of pendant edge lengths more stable than the other four approaches as the degree of interpolation rose (**Fig 2** rows 2-3). For analyses in the remainder of this section, we will use dates computed by the EQS-LS approach, but we have made distributions of trees available for both the EQS-LS algorithm and the birth model algorithm (see Data availability). A full breakdown of the results for each individual tree and algorithm is given in the supplementary material (S1 Fig, S2 Fig).

### Phylogenetic diversity in the complete tree

Our linear-time algorithms allowed us to estimate dates for all 2.3 million internal nodes of a complete bifurcating tree of extant described species, and therefore to estimate phylogenetic diversity for the entire tree and for any clade (**Fig 3**). We made use of phylogenetic diversity as a summary statistic to explore two dimensions of uncertainty around our trees.

**Fig 3.**
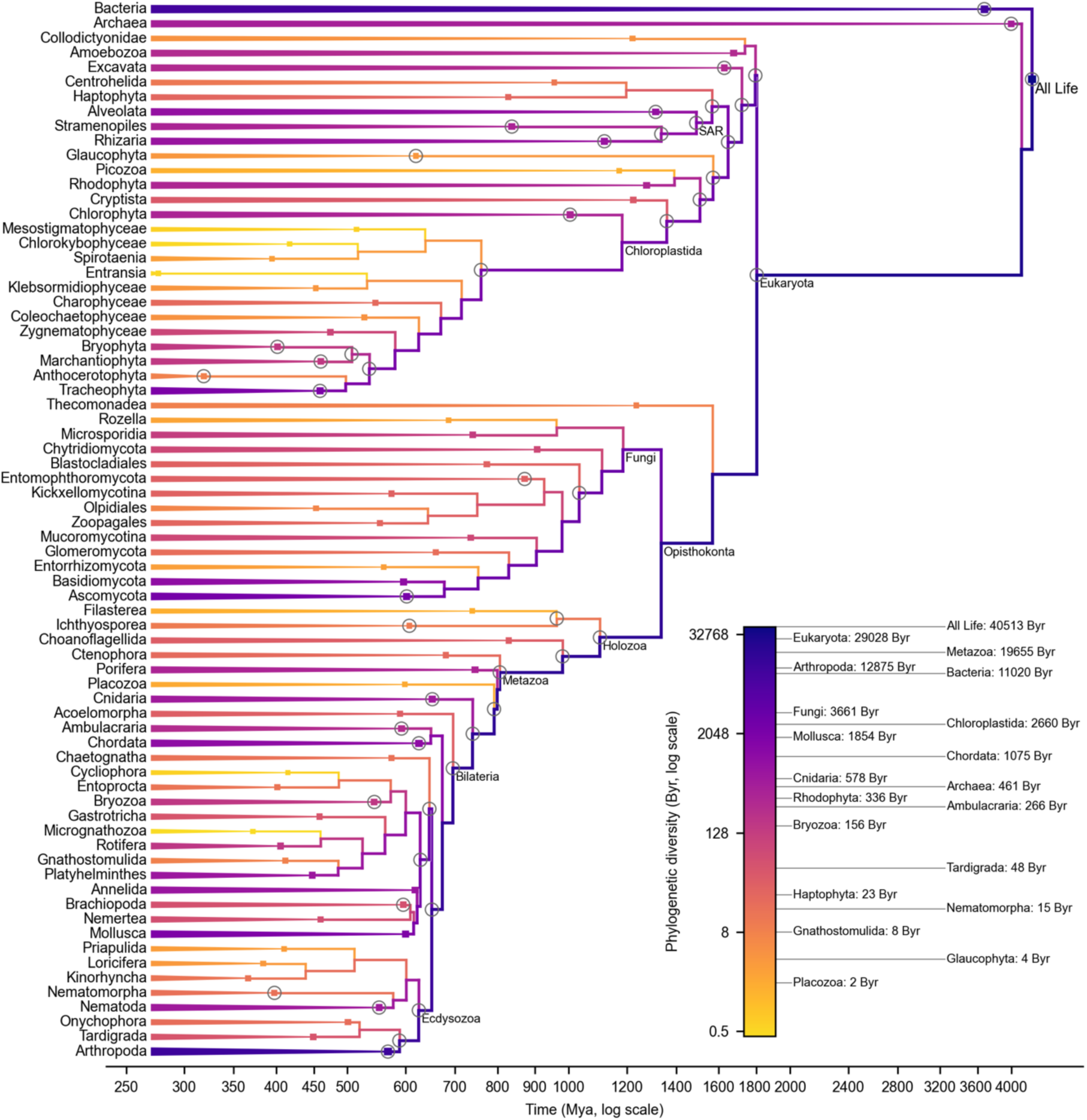
A pruned fully-dated complete phylogenetic tree of life from our distribution. The branch colours represent phylogenetic diversity (PD), illustrating how the PD of all life (upper right) is divided among clades in the tree. The underlying tree is ultrametric, but it is too large to display here. Instead we show a tree pruned to major clades: either infrakingdoms (where that rank exists in the Open Tree Taxonomy) or phyla, or the first taxon with rank lower than phylum if there is no phylum rootwards of it in the tree. The square nodes indicate the crown node and crown age of each clade. Circled nodes have calibration dates associated with them. PD estimates for selected clades of interest are given to the right of the colour bar; these estimates do not include the stem back to the origin of life.

First, we considered topological variation. We sampled 501 plausible topologies for the complete tree using our stochastic polytomy resolution algorithms (see Methods and materials). Where we had multiple sources for the date on a node, we used the median date. This produced a median phylogenetic diversity estimate for the complete tree of 39.6 trillion years (95% CI: 38.4-41.3 tyr; **Fig 4**). Highly-studied vertebrate and plant clades displayed very little topological variation (**Fig 5**), for example Chordata (98.5-101.4% of the median for the clade) and Tracheophyta (98.4%-101.6%). Most clades showed considerably wider variation (**Fig 5**), for example phylum Mollusca (82.5-116.1%) and phylum Rhodophyta (84.7-127.3%).

**Fig 4:**
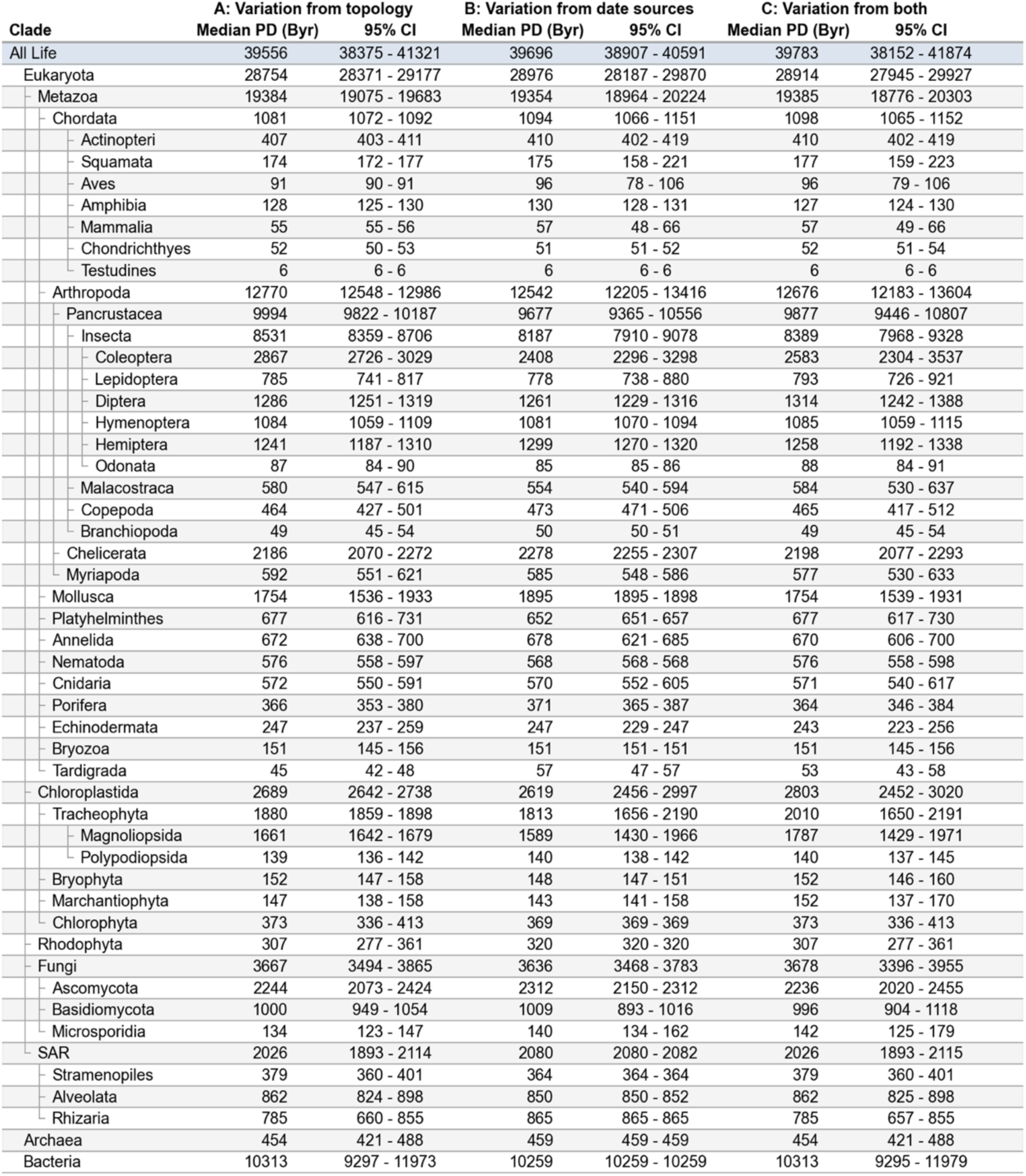
Table of phylogenetic diversity for selected clades of the tree. Panel A: PD owing to topological variation, from 501 different tree topologies using the median of the available dates for each node. Panel B: PD using a single tree topology, sampling 501 times from the available date sources. We used the topology corresponding to the median PD for the complete tree in the topological distribution. Panel C: PD from 1503 trees: 501 different topologies, with 3 sets of date source samples for each. In the hierarchy listed on the left, the clades chosen are a non-exhaustive subset of their parent clade.

**Fig 5.**
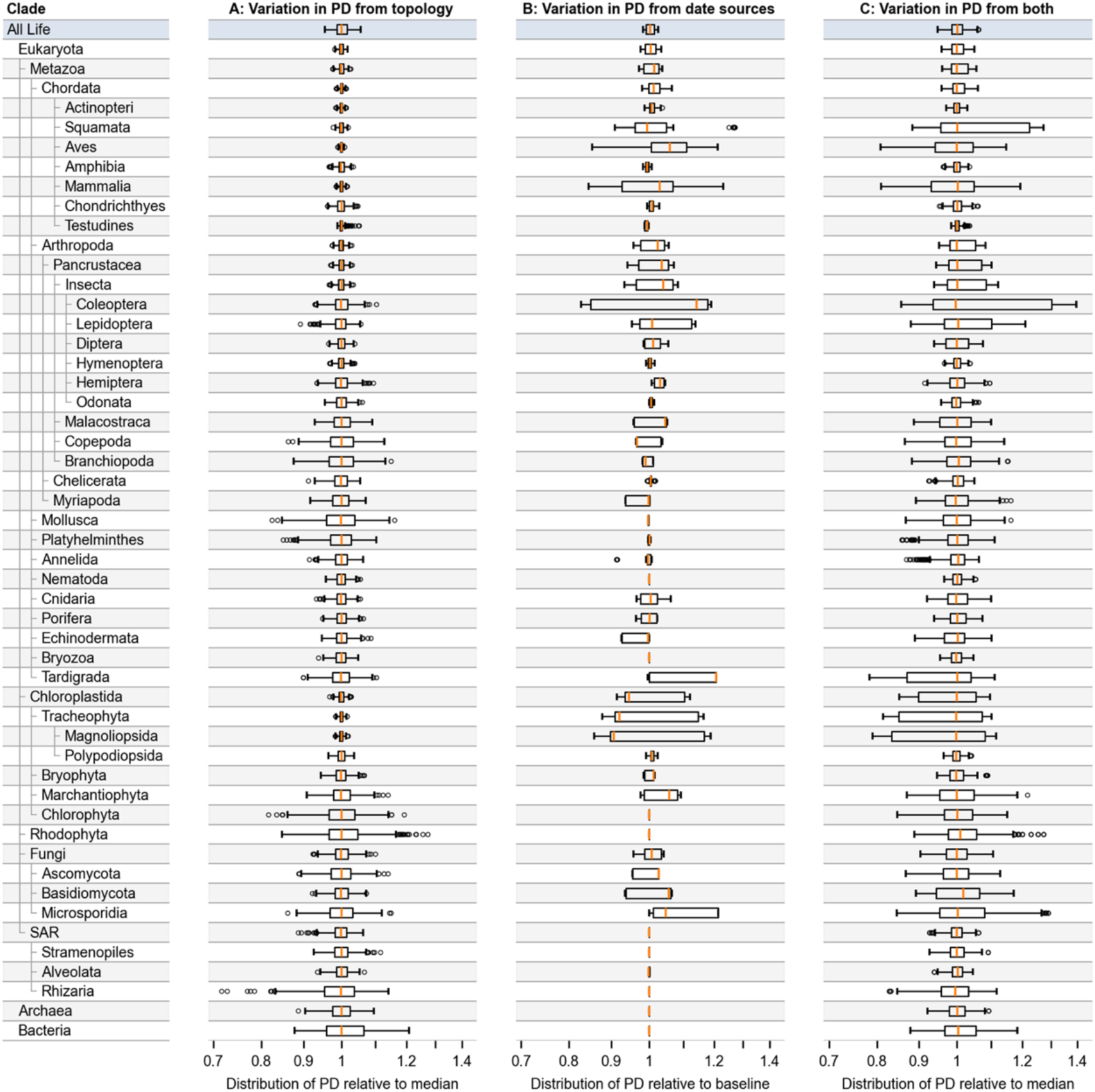
Phylogenetic diversity distributions for selected large clades of the tree. Panel A: distributions of PD owing to topological variation, from 501 different tree topologies using the median of the available dates for each node. Panel B: distributions of PD using a single tree topology, sampling 501 times from the available date sources. We used the topology corresponding to the median PD for the complete tree in the topological distribution. These PD distributions are shown relative to a “baseline” PD computed using median dates. Panel C: PD distributions from 1503 trees: 501 different topologies, with 3 sets of date source samples for each. In the hierarchy listed on the left, the clades chosen are a non-exhaustive subset of their parent clade.

Second, we considered temporal variation. Instead of using the median of the available date sources at each node, we randomly selected a single source (see Methods and materials). We repeated this 501 times, producing 501 different PD estimates for one tree with fixed topology. There was wide variation in PD in clades with a large number of sources such as birds (class Aves, 84.7-121.2% of the median for the clade) and mammals (class Mammalia, 84.5-123.0%) (**Fig 5**), while in nine clades there was no variation at all because no node in the clade had more than one date source (**Fig 5**).

Finally, we explored both dimensions of uncertainty simultaneously, sampling both topologies and date sources at random. We sampled 501 trees with three different sets of dates for each tree, giving a distribution of 1503 trees. This produced a median PD estimate of 39.8 trillion years (95% CI: 38.2-41.9 tyr; **Fig 4**).

## Discussion

### Fully-dated complete trees

The linear time complexity of our algorithms allowed us to resolve and date trees containing millions of species (**Fig 1**), which to the best of our knowledge has not previously been computationally feasible. Our trees incorporate all 2.3 million extant species in the Open Tree Taxonomy [25], which aims to include all formally-described species across all domains of life [41]. A fully-dated complete tree will be made available in the OneZoom tree viewer, which will allow users to explore the entire tree interactively [42].

To our knowledge, the largest fully-dated tree previously published is the TimeTree of Kumar et al. [43], an ongoing project that has the objective of including every species for which molecular data is available, and whose most recent published version in 2022 covered 137,306 species [44], about 6% of formally-described species. As of 2025, NCBI’s GenBank database contained at least some genetic information for 581,000 described species [45], a number that has increased by 15% over the past year [45]. The ongoing growth in data availability is encouraging, but it is clear that a significant proportion of species will remain unsequenced for the foreseeable future [46]. Our objective in this work was to give researchers access to a distribution of time-calibrated trees for the entire tree of life or any desired subset. We argue that efficient imputation and interpolation algorithms capable of filling in gaps in molecular coverage are, and will remain, an important tool in this endeavour, though we recognise that the resulting trees will not be suitable for every purpose [47–49].

### Interpolation quality

For our test trees or subtrees where at least 30% of nodes had input dates assigned to them, results for all five interpolation algorithms were similar (**Fig 2**). In this case, there is sufficient phylogenetic and temporal structure in the synthetic tree that the scope of interpolation error is relatively limited. In practice, only the highly-studied vertebrate clades have such strong coverage in the Open Tree database, with ferns coming close (29%) and other vascular plants falling short (8%). Elsewhere, few clades have date coverage of more 2%. Our analyses suggest that our EQS-LS algorithm is able to produce a reasonable distribution of branch lengths in trees with 5-10% of nodes dated (**Fig 2**), which represents a less daunting target than 30%. We hope that future researchers in under-represented clades will be able to help improve coverage to this level, by making their time-calibrated trees available electronically or contributing them directly to the Open Tree project.

### Phylogenetic diversity

Previous studies of phylogenetic diversity have mainly focused on vertebrate animals, and in particular mammals (class Mammalia) and birds (class Aves). For mammals, our median estimate of PD was 57 billion years, which is within the range of 37-66 billion years found across five previous studies [20, 50–53]. In birds, our median estimate is 96 billion years, broadly in line with figures of 74-90 billion years in four prior studies [50, 52–54]. A full list by clade of comparisons to previous studies is provided in the supporting information (S1 Table).

To our knowledge, two other studies have estimated PD across the entire tree of life. The review by Diaz and Malhi [52] cited PD estimates made by one of us (Rosindell) using a heuristic based on approximate clade age and species richness. In prokaryotes, Diaz and Malhi estimated PD of 221 trillion years across the two domains of Bacteria and Archaea, based on an estimated 740,000 operational taxonomic units (OTUs) [55–57]. Our tree instead contains only around 79,000 “species” of bacteria and archaea, as labelled in the Open Tree Taxonomy, resulting in the much smaller PD estimate from our tree of 10 trillion years (**Fig 5**). For eukaryotes, the PD estimate in Diaz and Malhi is 124 trillion years, based on an assumption of log-linear growth in species numbers over time. This assumption does not account for extinctions and will therefore tend to overestimate the ages of common ancestors, resulting in longer terminal branches and higher PD estimates. This explains why our PD estimate for eukaryotes of 29 trillion years is low by comparison. The second study to include a PD estimate for the entire tree, by Guo et al. [53], did so by computing species-level evolutionary distinctiveness (ED) scores [20], which can be summed to find PD for a clade. The ED scores were found by interpolating branch lengths along the path from each individual species to the tree root [53], without computing a single consistently-dated tree. Summing the scores across all species, this method found the total PD of all life to be 31 trillion years, compared to our median estimate of 39.8 trillion years with our EQS-LS interpolation algorithm. We applied the ED interpolation method to our fully-resolved tree, finding a median PD of 38.2 trillion years. This was very similar to our estimate, which indicates that the original difference resulted less from the method and more from the differing tree topologies. The prior study used the tree from the OneZoom tree viewer [42], which uses the Open Tree synthetic tree as a starting point but replaces certain clades of the tree with curated alternatives and resolves all polytomies purely at random [42]. We used only the base Open Tree and, where possible, resolve polytomies by imputing taxonomy-derived groups into the phylogenetic backbone, producing a different topology.

### Sources of uncertainty

In clades with few calibration dates, the choice of date interpolation algorithm will have a major effect on PD. Algorithms that create longer pendant edges will produce larger PD estimates, and vice versa. Here, we have used the EQS-LS algorithm because it produced the closest reproductions of dates and pendant edge lengths in the trees we tested (**Fig 2**). A distribution of trees produced using our alternative birth model interpolation algorithm is also available (see Data availability).

Having chosen the EQS-LS algorithm, our estimates of phylogenetic diversity incorporated two dimensions of uncertainty. First, we explored topological uncertainty: the distribution of plausible topologies for the tree of life created by the stochastic resolution of polytomies in the Open Tree. In the Open Tree synthesis algorithm, polytomies are created when taxonomic data is attached to the phylogenetic backbone. In clades with high phylogenetic coverage such as the vertebrates, there are few polytomies to resolve, and so we saw very low topological variation (**Fig 5**). In clades that come mainly from taxonomic data, such as molluscs (phylum Mollusca) and red algae (phylum Rhodophyta), we saw much broader distributions (**Fig 5**).

Second, we considered the temporal uncertainty resulting from variation across the available input dates. There are 76,935 nodes in the synthetic tree for which at least one published study has both estimated a date and been included in the Open Tree database. The majority of these nodes are in vertebrate animals and vascular plants. Here, nodes frequently had dates from multiple sources, and we were able to sample from the range of sources to reflect uncertainty (**Fig 5**), or use a median to summarise the information in multiple studies. Outside of these clades, to our knowledge most nodes in the tree of life have not been dated in any published study. For the small proportion of nodes where we did have a date, we usually had only a single source, and therefore we were not able to reflect uncertainty through sampling. This is a limitation in our inference of temporal uncertainty: if data are lacking this should imply higher uncertainty, not lower. Future work will include searching for studies not yet included in the Open Tree database, focusing on clades with low date coverage. Many source trees also included information on the uncertainty around their date estimates, but this data is not currently available in the Open Tree database.

There are further sources of uncertainty that could be considered in future. The crown ages of large, ancient clades where the descendant tree structure is derived largely from taxonomy are of particular importance to our method, because the crown ages will determine the dates interpolated across the entire clade. For example, the ages of the common ancestor of all Bacteria and of the largest fungus phylum Ascomycota are likely to have a significant effect on PD estimates for the complete tree, but we currently have only a single source for both these dates, so our analysis does not reflect this sensitivity.

We have also treated the set of extant species in the Open Tree Taxonomy as a fixed input. While the taxonomy attempts to incorporate all formally-described species, more species remain undescribed than have been described [58, 59]. Further, those undescribed species are likely to be concentrated in certain clades of the tree, such as Arthropoda [58, 60] and Fungi [61]. The incorporation of these “missing” species into the tree would alter the topology of some regions, and the interpolated dates and downstream analyses, in ways not reflected in our topological distributions. A PD estimate for the Arthropoda phylum that truly included every arthropod species on Earth would in all likelihood lie well beyond the bounds of our distribution, whereas our PD distribution for Chordata is likely closer to the truth.

Finally, we emphasise that our work summarises information from studies included in Open Tree’s database, but this does not include all published data. We are aware of dated trees not yet included in Open Tree, and where we are able to gain access to these trees in electronic form we plan to curate them into the Open Tree database, reducing uncertainty by expanding phylogenetic coverage and increasing the number of date calibration nodes. We are grateful to researchers who makes trees available electronically or inform us of regions of the tree that do not conform with the latest published work; indeed, we hope that researchers will consider contributing trees directly to the Open Tree project, to improve the resources available for all.

## Conclusion

We have designed and implemented algorithms to produce plausible candidates for the topology of the bifurcating tree of all life, and to interpolate dates across the tree, allowing us to produce fully-dated complete trees. We have used the trees to compute a distribution of phylogenetic diversity estimates for the whole tree of life. Our set of trees, as well as our polytomy resolution and date interpolation algorithms, are freely available (see Data availability); the input data we used and the seeds for the random number generator are included, making the trees reproducible.

Our work has built on the Open Tree of Life, the most complete tree presently available, deriving bifurcating trees that we think will be useful to new audiences. The source studies and algorithms used by Open Tree and Chronosynth are made available under the FAIR principles [62], and new time-calibrated trees can be curated into the Open Tree database by anyone with a GitHub account [23, 63]. The synthetic tree is regularly updated to incorporate newly-added phylogenetic and taxonomic information, and the computational efficiency of our algorithms will allow our distribution of trees to be updated in sync. We hope this work can spark new interest from potential contributors, who can further improve the Open Tree project for the benefit of the whole community, as well as enabling novel research that can take advantage of fully-dated complete trees.

## Materials and methods

### Overview of methods

We based our work on the Open Tree Synthetic Tree version 16.1 [27], which incorporates

2.3 million species, and the Open Tree Taxonomy 3.7.3 [64], which contains additional metadata about each taxon. We first pruned the tree to remove subspecies and extinct taxa, so that every leaf node represented an extant species. We then stochastically resolved the topology of the tree to produce a distribution of bifurcating trees, each one of which represented a plausible hypothesis for the evolutionary history of life. Finally, we mapped dates from the trees in the Open Tree database to the internal nodes in our trees [23, 33], and designed scalable algorithms to interpolate dates across all remaining undated internal nodes. These steps are summarised in **Fig 6**.

**Fig 6.**
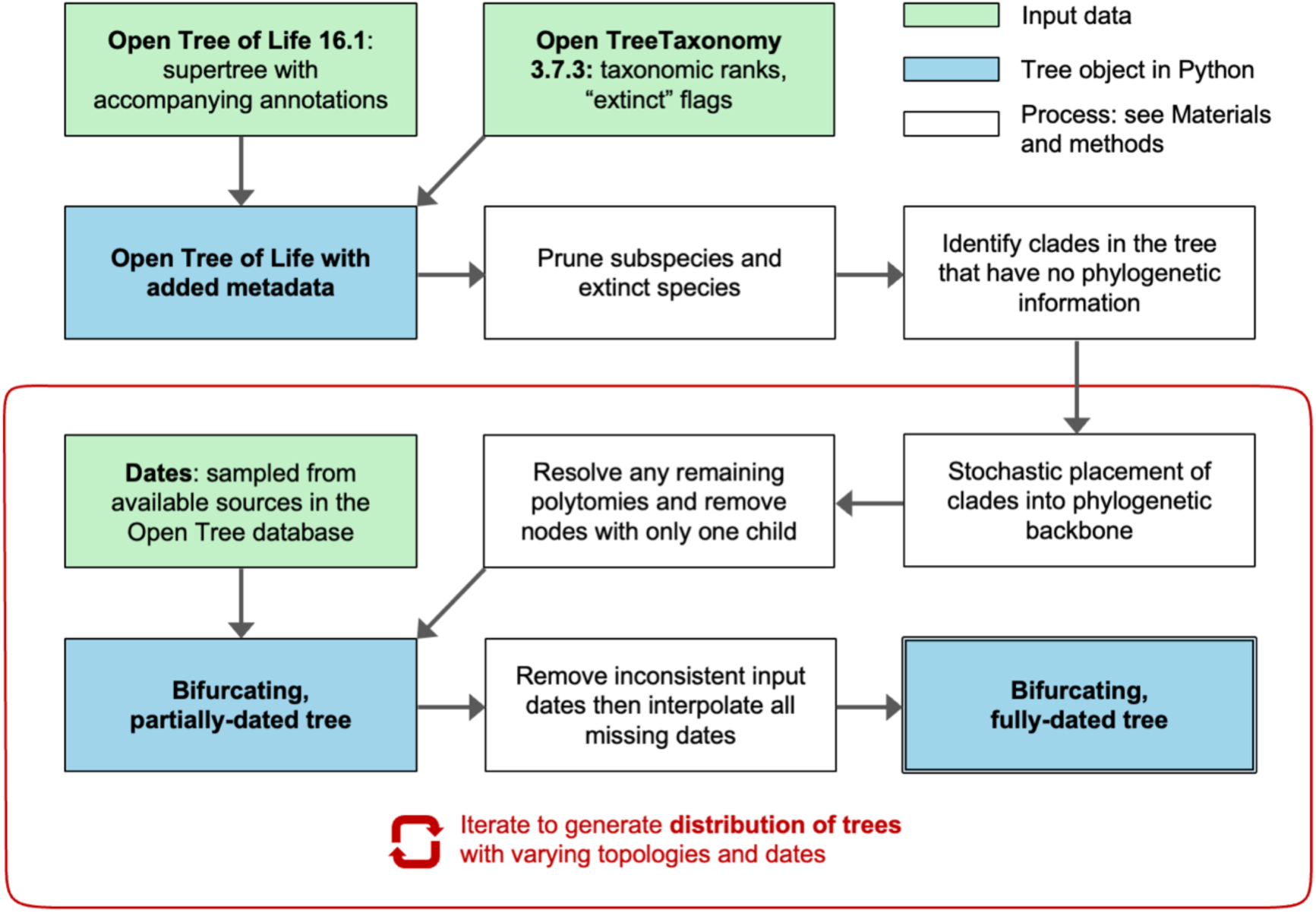
Summary of our methods for producing a distribution of dated trees. The iterative process shown will generate a distribution of trees incorporating both topological and temporal uncertainty. To explore only topological uncertainty, we used the median of available date sources at each node instead of sampling a new source with each iteration. To explore only temporal uncertainty, we used a single fixed topology instead of sampling a new topology with each iteration. Green boxes represent data inputs; white boxes summarise algorithmic processes detailed in the Materials and methods section of this paper; and blue boxes represent intermediate or final outputs.

### Initial pruning of the tree

Beginning with the labelled supertree (labelled_supertree_ottnames.tre) from the Open Tree of Life version 16.1 [27], we pruned any below-species-level nodes from the tree, including subspecies, varieties, formae, and leaf nodes labelled “no rank - terminal” in the taxonomy. If below-species nodes were present without a parent node representing their species, we kept one of them at random as a representative of that species. We removed nodes whose name contained the strings “uncultured” or “unidentified”, and removed 15 nodes whose name contained the string “intergeneric hybrids”. We pruned the birds clade (class Aves) to include only species present in the synthetic tree in

McTavish et al. [6], which is aligned to the Clements taxonomy [65], avoiding known issues with this clade in the Open Tree Taxonomy [6]. Similarly, we pruned the order of turtles (Testudines) to include only species present in the Turtles of the World checklist [66]. We removed the “extinct” flag that had erroneously been added to the species *Homo sapiens* in the latest version of the Open Tree Taxonomy. Finally, we removed any nodes labelled as extinct in the taxonomy and removed any of their ancestral nodes which may not themselves have been labelled extinct but had only extinct nodes below them.

These steps pruned 84,990 leaf nodes and 22,502 of their ancestors from the Open Tree. In addition, there were 1,718 leaf nodes that represented ranks above species, primarily empty genera; for each of these, we imputed a child node with the name “[Genus name]_sp._ott0000” as an unnamed representative species. The resulting tree had 2,294,776 leaf nodes, all now representing extant species, and 315,958 internal nodes.

### Topology

Our aim was to create a set of trees that were plausible hypotheses for the common descent of all life: that is, trees that respected the taxonomic hierarchy and were fully bifurcating. The steps to achieve this are described here, and illustrated using a small example tree (**Fig 7**).

**Fig 7.**
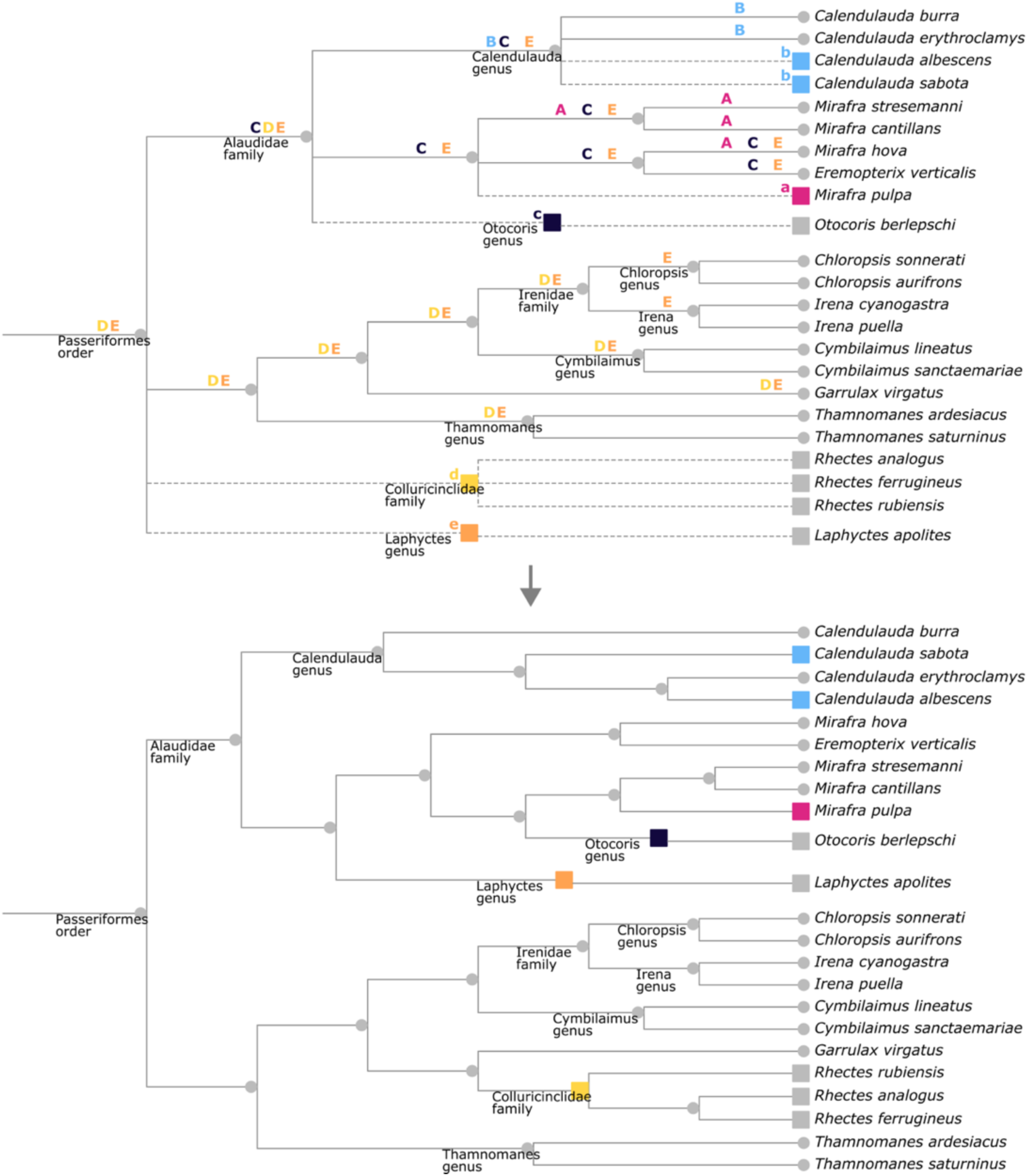
Illustration of our topology resolution algorithms. Top: A small phylogenetic tree illustrating how nodes in the Open Tree supertree that were absent from any underlying tree (squares, labelled with coloured lower case letters, with dashed-line branches above) were imputed into the phylogenetically-derived backbone (grey circles, with solid-line branches above). In random order, each coloured square node will be inserted into one of the branches indicated by the matching upper-case letters. For example, the *Laphyctes* genus is labelled with a lower-case “e”, and will be inserted into one of the branches labelled with an upper-case “E”, which are chosen to respect taxonomic hierarchy: they could be within any existing family in the backbone but not an existing genus. If the branch above the original parent node (Passeriformes in this example of *Laphyctes*) is chosen, the node to be moved would become sister to the entire descendant clade, with the original parent node remaining ancestral to both. Polytomy resolutions of this type were performed in random order, with newly-inserted groups added to the backbone as resolution progressed. As a final step, the polytomy in the *Rhectes* genus was resolved by choosing a topology at random. Bottom: one possible resolution, which will always be a bifurcating tree with taxonomic hierarchy respected and no existing monophyletic groups interrupted.

The Open Tree supertree algorithm first synthesises a bifurcating backbone tree comprising all the taxa included in their curated database of phylogenetic trees [23, 24]. Then, taxa in the Open Tree Taxonomy that are not present in any tree are connected to the crown node of their clade in the backbone, introducing polytomies. Working from our pruned tree, we used the information in the annotations.json file included with the Open Tree to label each node as derived either from the phylogenetic backbone or from the taxonomy. We then located all polytomies in the supertree and categorised them into three groups depending on their origin, resolving each category separately.

First, there were polytomies where taxonomy-derived groups had been attached to a phylogenetically-derived backbone node (this case is represented in **Fig 7** by the taxa labelled with lower-case b, c, d and e). There were 40,607 polytomous backbone nodes in this category. We resolved these using an approach similar to tree completion tools such as addTaxa [38], randtip [39], and pez [40], which insert taxonomy-derived species into randomly-chosen branches in a phylogenetic tree, applying certain location constraints. In our case, owing to the scale of the tree, location constraints were found algorithmically based on taxonomic information. For each polytomy, we detached the taxonomy-derived group, and searched below the polytomous node for candidate insertion points that would respect taxonomic hierarchy and would not interrupt any existing monophyly (for example, in **Fig 7**, candidate insertion points for the Colluricinclidae family are labelled with an upper-case D). We additionally included the possibility of the detached group becoming a sister to all the existing children of the backbone node. An insertion point was then selected randomly with uniform probability among the candidates. Successive resolutions of this type were performed in random order, with newly-inserted groups added to the backbone as resolution progressed.

Second, we resolved polytomies resulting from non-monophyletic genera in the supertree. In the phylogenetically-derived backbone tree, species from a genus can be placed into groups that also include other genera, if there was phylogenetic backing for those relationships. The supertree algorithm then takes taxonomy-derived species in the genus and attaches them to the most recent common ancestor of all their congeneric backbone species, creating a polytomy at that common ancestor. (This is illustrated in **Fig 7** by the Mirafra genus, where Mirafra pulpa has been attached to the common ancestor of the backbone Mirafra species). To resolve the 5,064 polytomies of this type we followed the approach of addTaxa [38]: we detached the taxonomy-derived species from the common ancestor, and imputed a new parent node for it into a branch in one of the congeneric monophyletic groups, choosing with uniform probability among all branches in all groups. (In **Fig 7**, the new location for M. pulpa is chosen uniformly among the branches labelled upper-case A.) As before, successive resolutions were performed in random order, with new branches added to the set of choices for potential insertion as resolution progressed.

Third, after resolving the previous two categories, there remained 74,402 polytomies where there was no phylogenetic information within a taxonomic group (as in the Rhectes genus in **Fig 7**). We resolved each of these by choosing a bifurcating topology uniformly at random from all possible topologies.

As a final step, we removed all unary nodes, which were created by the supertree algorithm only to provide taxonomic representation, producing a fully-bifurcating tree.

### Note on domains of life in the Open Tree

The current version of the Open Tree supertree has the root node of all life as the parent of the three domains Bacteria, Archaea, and Eukaryota. The presence of Bacteria and Archaea as sisters is derived from underlying phylogenetic trees, while Eukaryota is added from the taxonomy. We resolved this polytomy by making Eukaryota a sister to Archaea prior to any other polytomy resolution.

### Dating

#### Assigning dates to the trees

We used the Chronosynth library to assign dates from Open Tree’s underlying database of trees, called the phylesystem [23], to nodes in our bifurcating tree [6, 33]. Working with commit e060993 of the phylesystem (27 December 2025), we ran Chronosynth’s combine_ages_from_sources function, which searches the database for trees with branch lengths in millions of years, and then maps nodes in those source trees to nodes in the Open Tree synthetic tree by matching common ancestors [6].

We found 334 source trees (**Fig 8**; a full list of citations is given in S1 Text), and from these we were able to map at least one date to 76,953 nodes. The clade with the most source trees was birds (class Aves), with 129 sources in total and individual nodes having up to 44 sources (the crown nodes of Passeriformes and Galloanserae). In mammals (class Mammalia) there were 49 sources across the clade, with 15 date sources for the crown node of all mammals. Most dated nodes, however, had only one date source (48,938 nodes) or two (13,842 nodes) (S3 Fig).

**Fig 8:**
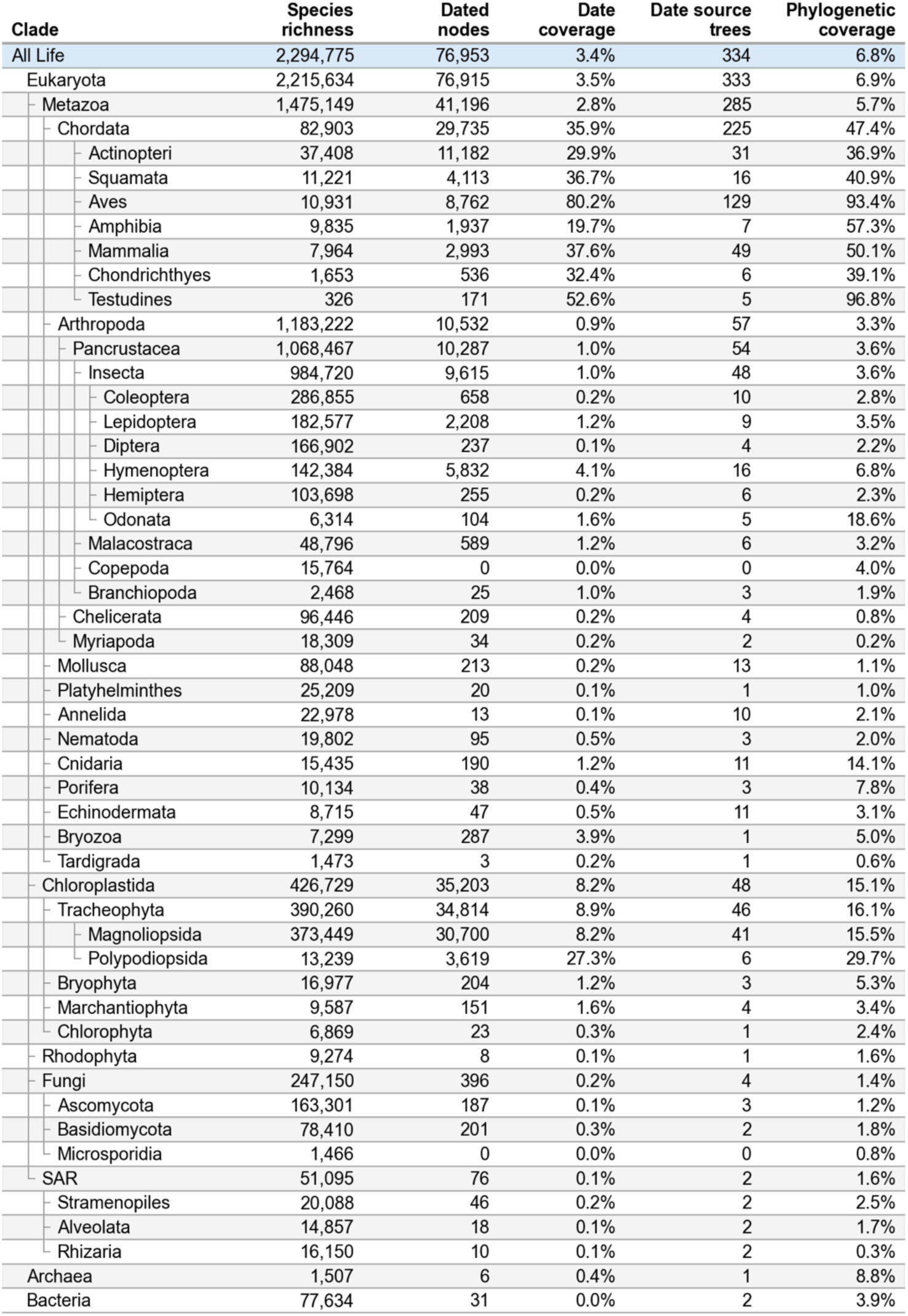
Breakdown of the calibration date coverage and phylogenetic coverage in our bifurcating tree. Date coverage is the percentage of internal nodes in the clade with at least one date source. Figures are for the original dates as mapped by Chronosynth, before checking for time consistency. In the hierarchy on the left, the clades chosen are a non-exhaustive subset of their parent clade.

Our date interpolation algorithms required a dated root node. We used a root date of 4247 million years ago (Mya), found in the TimeTree of Kumar et al. [44], who justified this date as lying between younger and older constraints on possible estimates for the origin of life. Their younger bound of >3970 Mya came from a molecular clock study of the divergence of archaea and eukaryotes [67], while the older bound of <4440 Mya was the earliest plausible cessation of asteroid impacts on the early Earth that evaporated the oceans and would have prevented life from developing [68].

### Algorithms for interpolating missing dates

Our goal was to interpolate a date on every undated internal node in a set of sparsely-dated trees with no branch lengths. We will refer here to “dated” and “undated” nodes, by which we mean internal nodes. All leaf nodes represented extant species and were assumed to have a date of zero. In describing our algorithms, we will use “down” to refer to movement from the root of the tree to its descendants toward the leaves, and “up” for the reverse movement from leaves to parents to ancestors toward the root.

We divided the interpolation into two distinct problems: interpolation between a dated node and one or more dated descendant internal nodes; and interpolation when a dated node had only undated internal nodes and leaf nodes below it. In the former case, paths from the dated node to each of its dated descendants must be considered to avoid negative branch lengths (see S4 Fig, and the discussion in the supplementary material of Kuhn et al. [28]). In the latter case, negative branch lengths can be avoided more straightforwardly by working down the tree and ensuring each undated node is assigned a date more recent than its parent. In our partially-dated bifurcating tree, 96.5% of undated nodes fell into the latter category.

We tested five interpolation algorithms, all of which we have implemented with run times that scale linearly with the number of nodes in a tree, making it feasible to use them across the entire tree of life. The first algorithm we will call equal splits or EQS, following Kuhn et al. [28], commonly used in the software Phylocom where its implementation is called branch length adjustment or BLADJ [34]. The algorithm assigns equal branch lengths to each branch along an undated path between a dated node and its oldest descendant. By identifying the oldest descendant, negative branch lengths are always avoided. For this reason, we applied only this algorithm to date the 3.5% of undated nodes located along undated paths between two dated internal nodes.

We compared the five interpolation algorithms across the remaining 96.5% of undated nodes, i.e. those located in clades where all internal nodes were undated except for a dated crown node. In these cases, there will be multiple undated paths covering the same time period (from the dated node to the leaves). For the EQS algorithm, the choice of which path is interpolated first will affect the result. In the BLADJ implementation, the path with the most nodes along it is dated first (**Fig 9**); we called this variant EQS-L, for “longest path”. We tested a second version of this where, instead of choosing the path with the most nodes, we first dated the path with the fewest nodes (**Fig 9**). We called this EQS-S, for “shortest path”.

**Fig 9:**
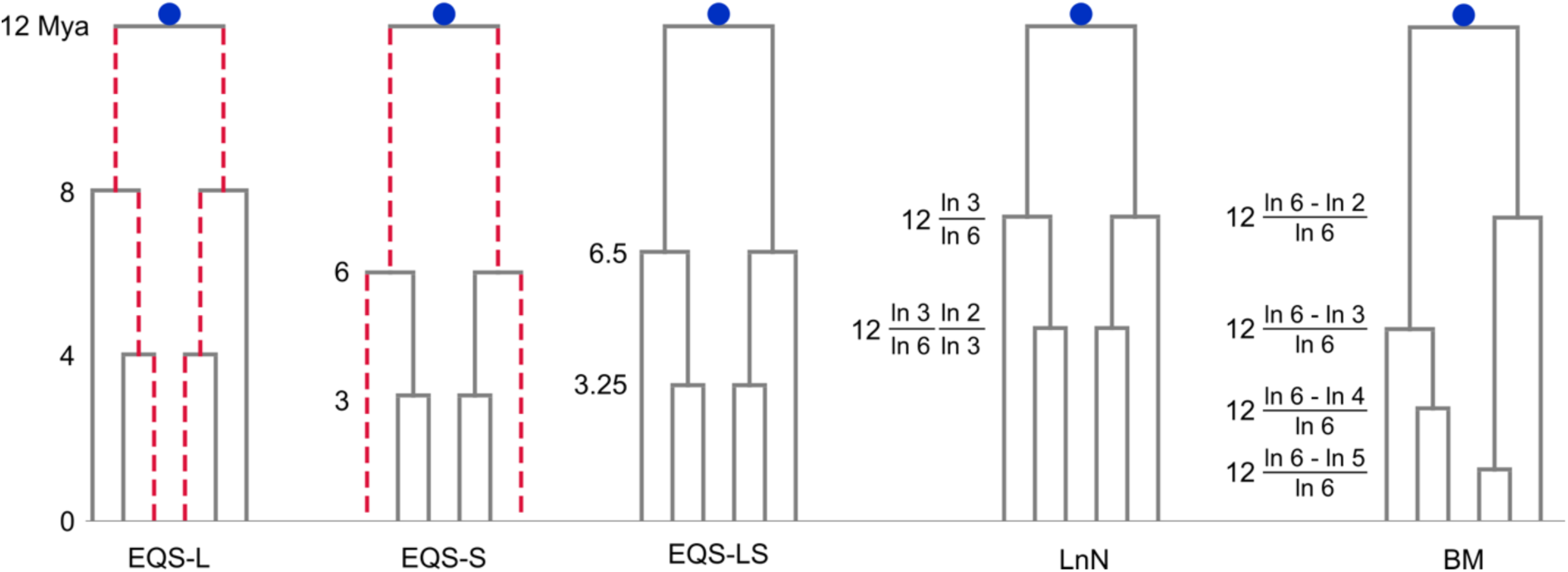
Examples of our five interpolation algorithms. The algorithms are illustrated in a clade where the only calibration date is 12 Mya on the crown node. The EQS-L and EQS-S algorithms space nodes evenly along undated paths, with the red dotted lines indicating the paths interpolated first - the longer path for EQS-L and the shorter path for EQS-S. The EQS-L algorithm produces pendant edges that are longer than or equal to the EQS-S algorithm. The EQS-LS algorithm computes dates as a linear combination of the two: 0.25*EQS-L + 0.75*EQS-S. The LnN and BM algorithms are inspired by birth models, with logarithmically-spaced dates. Here we have a crown age of 12 Mya, and a clade of six species. The LnN ages are (date of parent node) * (number of species in this clade) / (number of species in parent’s clade), applied recursively from crown to leaves down each lineage independently. The BM ages are (date of crown node) * (logarithm of the number of terminal lineages in the clade - logarithm of the number of lineages in the clade rootwards of this node in time) / (logarithm of the number of terminal lineages in the clade), so that the number of lineages through time in the clade plotted on a semilog plot is straight line.

Our third algorithm was a linear combination of the dates from these two approaches, inspired by observing that the EQS-L approach leads to very long pendant edges relative to a fully-dated tree, while the EQS-S approach produces extremely short pendant edges. To find a suitable combination, we used the four published trees described in the section below on quality assessment, which had molecular data for all species. We found that assigning dates as 0.25*(EQS-L date) + 0.75*(EQS-S date) minimised error relative to the published dates. We called this algorithm EQS-LS. Since both input algorithms produce a consistent set of dates across the tree, and the EQS-S dates must always be younger or equal to the EQS-L dates, any linear combination of the two must also produce a consistent set of dates.

The fourth algorithm we tested was proposed by Purvis and Nee [35], based on a birth or birth-death model where the age of a clade is in expectation proportional to the logarithm of the number of species in the clade. This method, which we called LnN following Kuhn et al. [28], computes a date for each undated child node by multiplying the date of its parent by the ratio of the logarithm of the species counts in the child clade and the parent clade. Given a dated root of the clade, this can be applied recursively down a tree to date all nodes (**Fig 9**).

These four algorithms are deterministic, i.e. given a tree topology and a set of dated nodes they will produce the same result on any run. Our fifth algorithm was, like the LnN approach, based on a birth model with exponentially-distributed waiting times, but added a stochastic element. Rather than working independently down each lineage of a clade as the LnN algorithm does, we instead considered all speciation events in the clade and assigned their time ordering at random: working down a clade, the probability of a lineage being the site of the next speciation event is proportional to that lineage’s fraction of the total remaining speciation events. The dates for the speciation events are chosen so that the number of lineages grows exponentially over time (**Fig 9**); that is, a lineages-through-time semilog plot would show a straight line. Specifically, the sequence of dates is (logarithm of the number of terminal lineages in the clade - logarithm of the number of lineages in the clade rootwards of this node in time) / (logarithm of the number of terminal lineages in the clade). We called this the BM algorithm, for birth model.

### Implementation and time complexity of date interpolation algorithms

We implemented all algorithms using a dynamic programming approach with run times that scaled linearly with the number of nodes (**Fig 1**). Here we describe the implementation of the EQS-L algorithm, and then summarise the modifications required by the other four algorithms.

The first step in the EQS-L algorithm was to traverse the tree and label each undated node with the information required to interpolate missing dates: we require the date of the oldest descendant of the node, and the number of nodes along the path to that date (for an illustrative example of the procedure described here, see **Fig 10**, top). We begin with leaf nodes, which we label with the tuple (0, 0), representing a date of 0 and a path length to that date of 0. These tuples are reported to each node’s parent. Then, if a parent is undated, we look at the tuples it received from its children, and label it with the tuple that contained the oldest date, incrementing the path length in the tuple by one to represent the additional node now in the path. If multiple tuples are received with the same date, the tuple with the longest path is retained (for example, in **Fig 10**, the common ancestor of species M and P stores the path length to M). If the parent is dated, it is labelled with a tuple containing its own date and a path length of 0. In either case, the parent reports its tuple to its own parent, and this process continues recursively up to the tree root.

**Fig 10.**
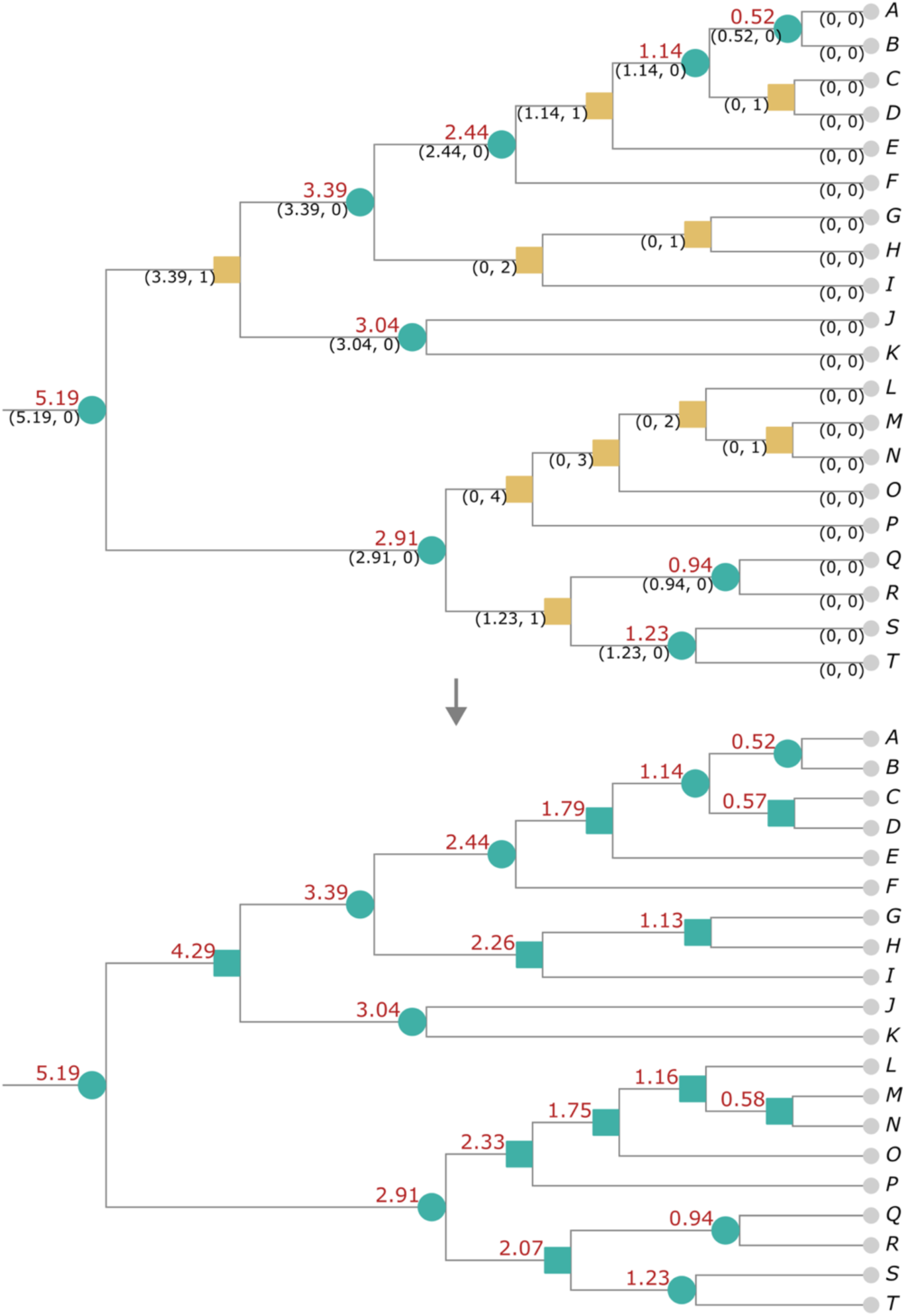
Illustration of the EQS-L algorithm for evenly interpolating missing dates between known dates. Top: Leaf nodes are shown as small grey circles, and are given a date of zero. Nodes with dates assigned are shown as blue-green circles, with the date given in red text above the branches. The yellow-brown squares are the nodes to be dated. Each node is shown with the tuple reported upwards, containing: (oldest date reported to this node, path length down the tree to that date). If the oldest date reported is a tie, the longer path length is used; for example, the common ancestor of M and P retains the path towards M. For the “shortest path” EQS-S version of the algorithm, the path to P would be retained. Bottom: Working down the tree from the dated root node, each undated node is given a date spaced evenly along the path between its parent and the oldest date below it, which is stored in its label.

Once all internal nodes are labelled we begin again, this time from the root, working down the tree and assigning a date to every undated node. Since we require a dated root node and work down the tree, a node’s parent can be assumed to be dated. We therefore space each undated node evenly along the path between the date of its parent and the oldest date beneath it, using the oldest date and path length information recorded in the node’s label (**Fig 10**, bottom). This completes the dating of the tree in linear time, using one postorder and one preorder traversal.

For the shortest path EQS-S algorithm, the procedure is the same except that when a node has received multiple tuples with the same date, it retains the tuple with the shortest path rather than the longest. The LnN algorithm uses similar recursive logic to first label each node with the number of leaf nodes below it. It then uses the oldest-descendant labels to identify when a dated node has only undated nodes and leaf nodes below it, and in those cases, works down from the root assigning a date to each undated child by multiplying the date of its parent node by the ratio of the logarithm of the leaf counts in the child clade and the parent clade. The BM algorithm also identifies dated nodes with only undated and leaf nodes below, then works down the clade, randomly choosing among the lineages and assigning the expected date for each bifurcation given the numbers of lineages.

### Quality assessment of date interpolation algorithms

We assessed the interpolation algorithms by removing input dates at random from a clade, interpolating all missing dates, and measuring the difference between the interpolated dates and the input dates that we had removed. We did this first for four published dated trees, each with molecular data for all species: the Jetz et al. tree of 6,645 bird species [69]; the tree of 4,988 frog species by Portik et al. [70]; the mammals tree of Upham et al. with 4,064 species [71]; and the tree of 609 cartilaginous fish by Stein et al. [72].

For each tree, we first removed a fixed proportion of dates - for example, 10% - at random from the internal nodes of the tree, though never from the root node. We ran the first interpolation algorithm over the tree, and computed the absolute difference in million years between the interpolated dates and the original dates. We then divided the differences by the root age of the tree, to normalise across trees of different ages - average absolute differences can be expected to be larger if all the dates in the tree are larger (older). We then ran the other four algorithms on the same partly-dated tree. That completed one replicate for each algorithm.

We then restored all input dates and randomly chose a different 10% of dates to remove, again finding the interpolation errors for each algorithm, and repeated this until we had 10 replicates for each algorithm. The errors were then averaged across the ten replicates. This gave us one datapoint for each algorithm: the average interpolation error when we removed 10% of dates from one fully-dated tree. We did this for all four trees, and averaged the errors. Then we removed 25% of dates, repeating the entire process, and so on; we tested removing 10%, 25%, 33%, 50%, 60%, 70%, 80%, 90%, 93%, 96%, 98%, 99%, and 100% of the internal dates (excluding the root age).

At the same time, we computed for each replicate the mean and median pendant edge length in the tree, relative to the mean and median in the original fully-dated tree. We considered it desirable for the distribution of pendant edge lengths to remain close to the original distribution when interpolating dates, i.e. for the relative edge lengths to stay close to 1.

Finally, we repeated this analysis on seven clades of our bifurcating complete tree, chosen because they had a high proportion of their nodes dated including the crown node of the clade. The clades were: Aves (10,931 spp., 73% dated); Squamata (11,221 spp., 34% dated); Mammalia (7964 spp., 33% dated); Actinopteri (37,408 spp., 28% dated); Polypodiales (10,709 spp., 27% dated); Amphibia (9,835 spp., 19% dated); and Fagales (2,274 spp., 20% dated). Interpolation errors were measured relative to the crown age of the clade. Since we had no fully-dated baseline tree, pendant edge lengths were normalised relative to those resulting from the EQS-LS algorithm when no dates had been removed.

### Date consistency across the tree

The dates we used came from 334 trees (a full list of citations is given in S1 Text). The dates and topologies in these trees were not necessarily consistent with one another. When the dates were assigned to the synthetic supertree, some nodes had dates younger than a descendant or older than an ancestor, which would imply negative branch lengths. The BLADJ implementation in Phylocom of the equal splits algorithm deals with inconsistent dates by working down the tree and removing any date older than an ancestral date [34]. This approach has linear time complexity, but by working in one direction from the root it will bias the results toward younger dates, and is likely to remove more dates than necessary to produce a consistent tree. For example, consider a node with a date of 10 Mya that has two older children dated 11 and 12 Mya. The BLADJ algorithm will remove the dates on the children because they are lower in the tree, but consistency could also be achieved by removing only the parent date of 10 Mya, retaining two of the original three dates rather than one.

We developed an alternative algorithm that removes the minimum number of dates necessary to make a tree consistent. We first traversed once through the tree and produced for each node a list of any older descendants and younger ancestors. Then we chose the node with the largest number of inconsistencies, removed the date from that node, then removed the node from the lists for all other nodes. We then chose the node with the most remaining inconsistencies, and so on until there were no inconsistencies left.

Our method has a worst-case time complexity of O(n × m), where n is the number of nodes and m is the number of dated nodes: we visit each node once (n nodes), and at each node we scan the list of all dated descendants (m nodes). On our sparsely-dated trees we have m << n, so the realised time complexity is very close to linear in the number of nodes (**Fig 1**).

### Phylogenetic diversity distributions

For each fully-dated tree, we summed all branch lengths to compute phylogenetic diversity. We traversed the tree from leaves to root; at each node we stored the sum of desendant branch lengths, then added the length of the branch to the parent node and passed this sum on to the parent. This recursion computes and stores the PD of the clade below every node in the tree in linear time.

We then computed three PD distributions: one exploring temporal uncertainty, one exploring topological uncertainty and one exploring both topological and temporal uncertainty simultaneously.

First, temporal uncertainty in the tree was explored by choosing among the available date sources at each node. Here, we used a single fixed tree topology; we chose the topology that give us the median PD estimate from the distribution exploring topological uncertainty. We created a list of all date source trees. We randomly shuffled the list, then treated this list as a priority ordering of sources: when we assigned dates to nodes in the tree, we used the date from the highest-priority source in the list. We then made the dates time-consistent, interpolated all dates and computed PD. Then we removed all the dates, re-shuffled the list of sources, and repeated the process 501 times, producing a distribution of 501 PD estimates.

Second, we investigated topological uncertainty by running the stochastic polytomy resolution step repeatedly to produce 501 different dated trees. For each tree topology generated, we computed a PD estimate using the median of the available date sources at each node. We then removed all dates and assigned a single date source at each node using random prioritisation of sources as previously described. We repeated this to give us three different dated trees with the same topology but varying dates. These steps produced two further distributions, one of 501 PD estimates exploring topological variation only, and one of 1503 PD estimates exploring both topological and temporal variation simultaneously.

### Benchmarking and software

We compared the run times of our EQS-L interpolation implementation and the BLADJ implementation in Phylocom [34] on increasingly large, partially-dated subtrees of our complete tree. The performance of BLADJ was measured using the software “hyperfine” [73] in the zsh shell on macOS Sequoia 15.1.1, running on an Apple M4 Pro Macbook Pro. The Phylocom software was compiled for Apple Silicon from source on the same machine. For benchmarking, our code read the same input files and wrote the same outputs as Phylocom, and was run in the CPython 3.13.0 implementation of Python using IPython’s “%timeit” magic command to measure run times. Random numbers were generated using the default random number generator in Numpy 2.1.3 [74]. For tree structures and tree visualisation we used the ETE4 library [75].

## Data availability

The Python code for all the algorithms described here is available at https://github.com/jdduke24/dated-complete-tree. Distributions of fully-dated complete trees using both the EQS-LS and birth model interpolation algorithms are available at https://doi.org/10.5281/zenodo.19049120. All input trees are available on Github in the Open Tree data store https://github.com/OpenTreeOfLife/phylesystem-1 and via the OpenTree data curation website https://tree.opentreeoflife.org/curator.

## Supporting information

**S1 Fig:**
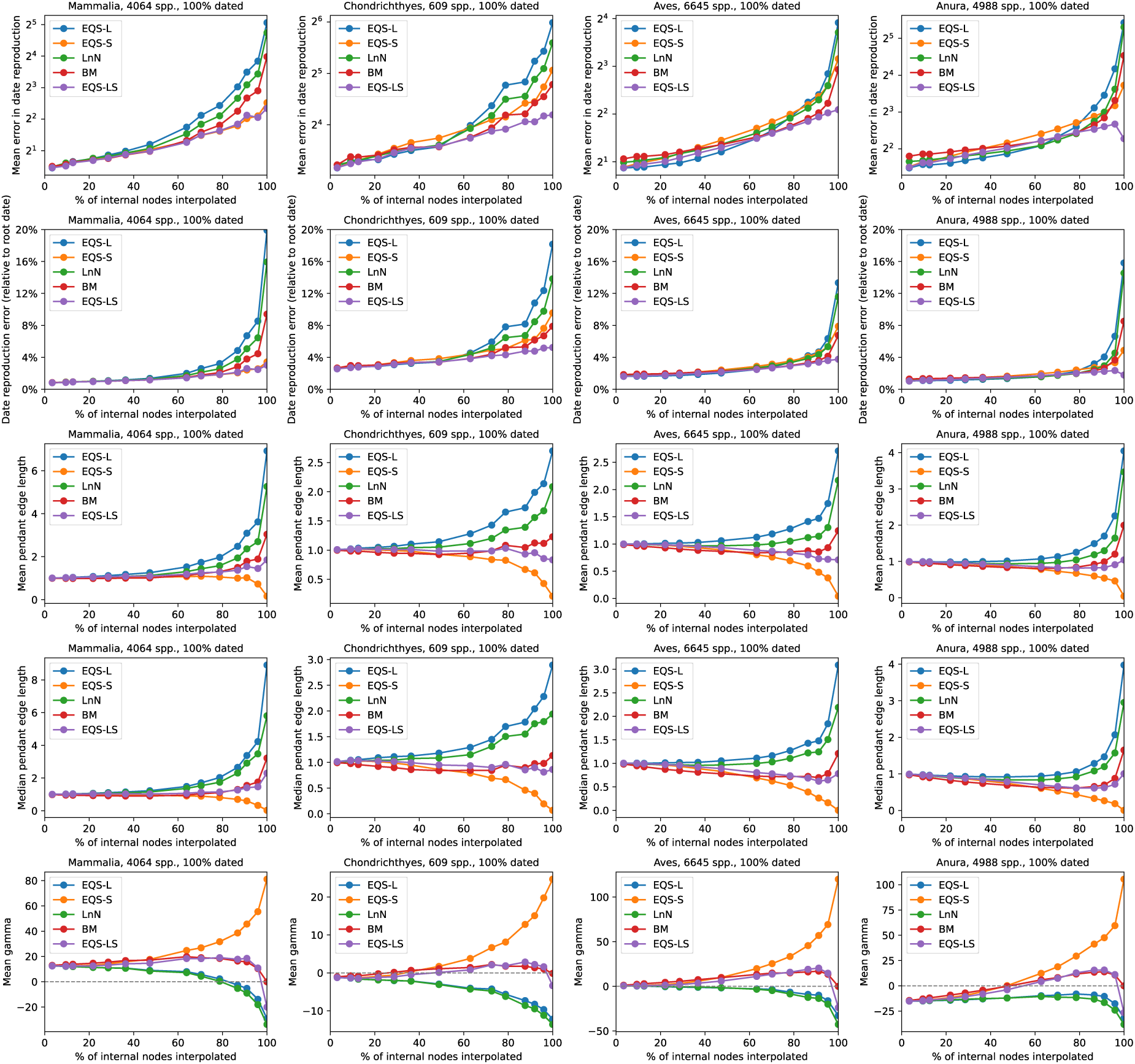
Full breakdown of date interpolation quality assessment on four fully-dated trees. Columns from left to right: results for four fully-dated trees with molecular data for all included species: the mammals tree of Upham et al. [71]; the tree of 609 cartilaginous fish by Stein et al. [72]; the tree of 4,988 frog species by Portik et al. [70]; and the Jetz et al. tree of 6,645 bird species [69]. We removed a proportion of dates at random from the tree (but never the root age), then interpolated dates across all undated internal nodes and compared trees with interpolated dates to the original trees. Top row: the average error in date reproduction tests (absolute difference in million years). Second row: the average error in date reproduction as a proportion of the root age of the tree. This normalises across trees of different ages; for example, sharks are older than mammals, so the absolute differences are expected to be larger. Third row: the mean pendant edge lengths for the same trees, divided by the mean for the fully-dated tree. Fourth row: Median pendant edge lengths, divided by the median for the fully-dated tree. Fifth row: Pybus and Harvey’s gamma statistic characterising the distribution of branch lengths [76]. Positive values indicate younger dates and shorter pendant edges than expected from a birth model, and vice versa. All trees converge on 0 when 100% of dates are interpolated using birth model interpolation (red lines), as expected.

**S2 Fig:**
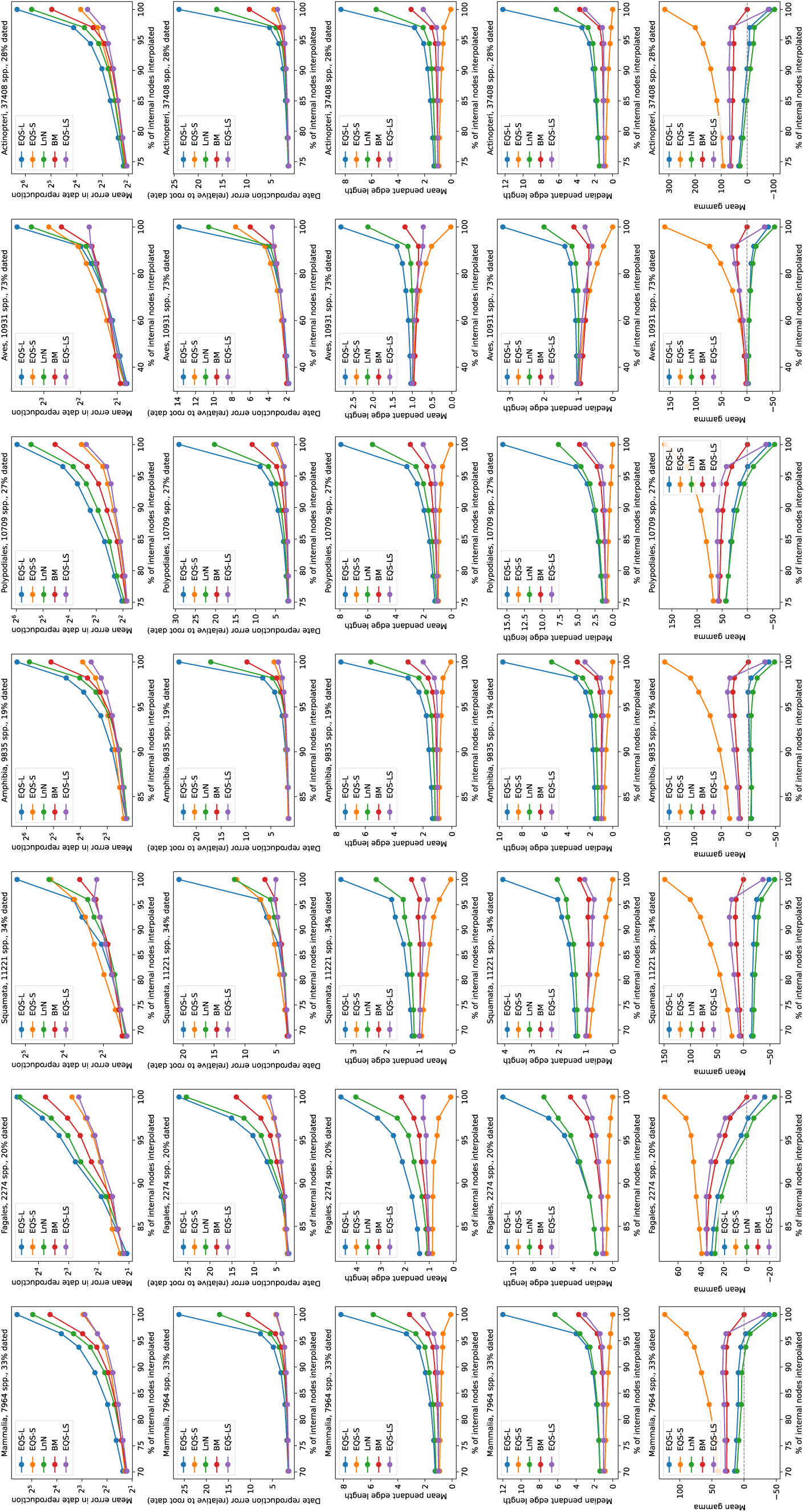
As in S1 Fig but for seven subtrees of a partially-dated complete tree. Note the different x and y scales in each column.

**S3 Fig:**
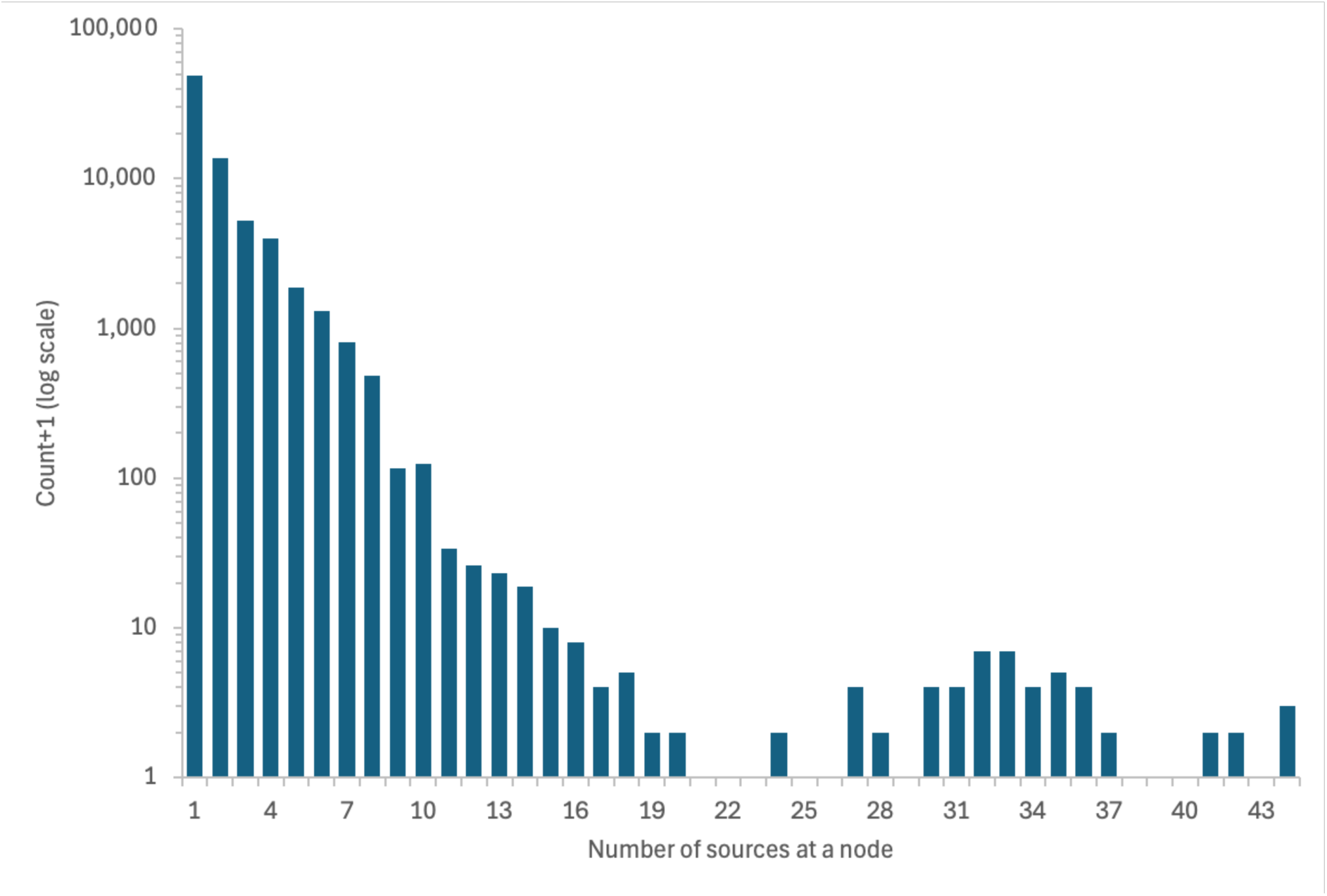
Distribution of the number of date sources at nodes with at least one source. There were 76,935 nodes with at least one date source. Around 82% (62,780) of these dated nodes had only one or two sources; 99% of nodes have seven or fewer sources. There are 25 nodes with 20 or more sources, all in the birds clade.

**S4 Fig:**
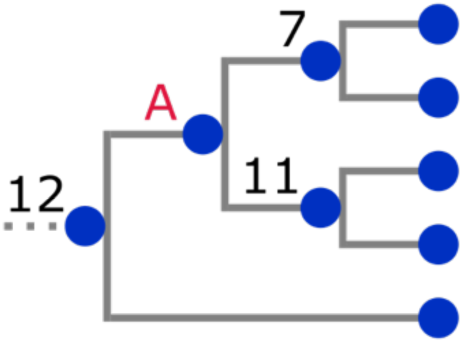
Interpolation can result in negative branch lengths. The numbers shown next to the internal nodes in this cladogram are dates in Mya; the node labelled A is to be dated. Assigning a date to A using, for example, 12 x ln(4) / ln(5) = 10.3 Mya, as computed by the LnN algorithm explained in the main text, would result in a negative branch length between A and the node dated 11. The EQS algorithms find the oldest dated node leafwards of A, and place A midway in time between the node dated 12 and the oldest dated node below it (11), guaranteeing no negative branch lengths.

**S1 Table:** Comparisons of our median phylogenetic diversity estimates with estimates from previous studies.

| Clade | Study | Species richness | PD (Myr) | PD:spp ratio |
| --- | --- | --- | --- | --- |
| **All Life** | This study | 2,294,776 | 39,782,514 | 17.3 |
|  | Guo et al. (2025) [50] | 2,235,076 | 31,000,000 | 13.9 |
|  | Diaz & Malhi (2022) [52] | 2,850,746 | 345,367,000 | 121.1 |
| **Eukaryotes** | This study | 2,215,634 | 28,914,060 | 13.1 |
| (Eukaryota) | Guo et al. (2025) | 2,174,404 | 26,004,213 | 12.0 |
|  | Diaz & Malhi (2022) | 2,110,866 | 124,417,000 | 58.9 |
| **Mammals** | This study | 7,965 | 56,828 | 7.1 |
| (Mammalia) | Gumbs et al. (2024) | 6,253 | 37,068 | 5.9 |
|  | Martyn et al. (2012) [51] | 5,139 | 49,131 | 9.6 |
|  | Isaac et al. (2007) [20] | 4,433 | 42,628 | 9.6 |
|  | Guo et al. (2025) | 5,046 | 50,075 | 9.9 |
|  | Diaz & Malhi (2022) | 5,045 | 66,000 | 13.1 |
| **Birds** | This study | 10,931 | 96,272 | 8.8 |
| (Aves) | Gumbs et al. (2024) | 10,988 | 89,747 | 8.2 |
|  | Jetz et al. (2014) [54] | 9,993 | 77,149 | 7.7 |
|  | Guo et al. (2025) | 10,002 | 74,177 | 7.4 |
|  | Diaz & Malhi (2022) | 9,993 | 81,000 | 8.1 |
| **Ray-finned fish** | This study | 37,408 | 410,443 | 11.0 |
| (Actinopterygii) | Gumbs et al. (2024) | 32,760 | 329,935 | 10.1 |
|  | Guo et al. (2025) | 34,432 | 348,288 | 10.1 |
|  | Diaz & Malhi (2022) | 35,810 | 1,512,000 | 42.2 |
| **Amphibians** | This study | 9,835 | 126,956 | 12.9 |
| (Amphibia) | Gumbs et al. (2024) | 8,024 | 142,670 | 17.8 |
|  | Isaac et al. (2012) | 4,331 | 57,980 | 13.4 |
|  | Guo et al. (2025) | 8,688 | 95,735 | 11.0 |
| **Lepidosaurs** | This study | 11,221 | 176,652 | 15.7 |
| (Lepidosauria) | Gumbs et al. (2024) | 10,735 | 134,253 | 12.5 |
|  | Tonini et al. (2016) [77] | 9,755 | 122,478 | 12.6 |
| **Gymnosperms** | This study | 2,014 | 38,360 | 19.0 |
| (Acrogymnospermae) | Forest et al. (2018) [78] | 1,090 | 12,671 | 11.6 |
| **Jawed vertebrates** | This study | 79,430 | 895,692 | 11.3 |
| (Gnathostomata) | Gumbs et al. (2024) | 70,426 | 782,102 | 11.1 |
| **Cartilaginous fish** | This study | 1,653 | 52,119 | 31.5 |
| (Chondrichthyes) | Gumbs et al. (2024) | 1,290 | 41,297 | 32.0 |
| **Crocodylians** | This study | 34 | 586 | 17.2 |
| (Crocodylia) | Gumbs et al. (2024) | 25 | 613 | 24.5 |
| **Turtles** | This study | 326 | 6,245 | 19.2 |
| (Testudines) | Gumbs et al. (2024) | 351 | 6,520 | 18.6 |

**S1 Text: References for studies in the Open Tree phylesystem that contain dated phylogenetic trees used as date sources in this work.** Some studies contributed more than one tree. Listed in alphabetical order of first author surname.

